# Biofilm Formation and Virulence of *Shigella flexneri* is Modulated by pH of Gastrointestinal Tract

**DOI:** 10.1101/2020.10.16.336651

**Authors:** I-Ling Chiang, Yi Wang, Satoru Fujii, Brian D. Muegge, Qiuhe Lu, Thaddeus Stappenbeck

## Abstract

*Shigella* infections remain a major public health issue in developing countries. One model of *Shigella* pathogenesis suggests that the microfold epithelial cells in the small intestine are the preferred initial site of infection. However, a growing body of evidence supports an alternative model whereby *Shigella* primarily infects a much wider range of epithelial cells that reside primarily within the colon. Here, we investigated whether the luminal pH difference between the small intestine and colon could provide evidence in support of either model of *Shigella flexneri* pathogenesis. As virulence factors leading to cellular invasion are linked to biofilms in *S. flexneri*, we examined the effect of pH on *S. flexneri*’s ability to form and maintain adherent biofilms when induced by deoxycholate. We showed that a basic pH inhibited formation and dispersed pre-assembled mature biofilms while an acidic pH (similar to the colonic environment) did not have either of these effects. To further elucidate this phenomenon at the molecular level, we probed the transcriptomes of biofilms and *S. flexneri* grown in different pH conditions. We identified specific amino acid metabolic pathways (cysteine and arginine) that were enriched in the bacteria that formed the biofilms, but decreased upon pH increase. We then utilized a type III secretion system reporter strain to show that increasing pH reduced deoxycholate-induced virulence of *S. flexneri* in a dose dependent manner. Taken together, these experiments support a model whereby *Shigella* infection is favored in the colon because of the local pH differences in these organs.

## Introduction

*Shigella* spp. are gram-negative bacterial pathogens that cause approximately 5-15% of the diarrheal burden in the world (1, 2). Infection can have serious consequences as *Shigella* infection results in ~200,000 deaths per year, mostly in pediatric populations of developing countries (2). *Shigella* is most efficiently transmitted human to human by a fecal-oral route and can invade intestinal epithelial cells (3, 4). Despite the morbidity and mortality resulting from *Shigella* infections, clinically termed shigellosis, there is not an effective vaccine for the bacteria nor do we fully understand the mechanism by which this pathogen infects the gastrointestinal tract (5).

One major limitation that has slowed our understanding of *Shigella* infections is that humans are the main reservoir for this microbe. Primates can be infected, but establishment of other *in vivo* animal infection models that recapitulate the phenotypes of enteric invasion has been challenging (6). Data from infection of *ex vivo* ligated rabbit ileal loops suggests that *Shigella* are phagocytosed by specialized epithelial microfold (“M”) cells that are most prominent overlying organized gut-associated lymphoid tissue (GALT). These M cells facilitate bacterial translocation to immune cells that underlie their basolateral surface (7). One shortcoming of this model is that in humans M cells are primarily found in ileal Peyer’s patches of the small intestine, whereas *Shigella* infection occurs predominantly in the colon (1, 8). Physiological and pathological studies of *Shigella* infections in humans and rhesus monkeys, respectively, demonstrate that the bacteria primarily invade and damage the colonic epithelium in areas not necessarily associated with GALT and has minimal effects on the small intestine (9, 10). However, *in vitro* primary gut epithelium infection studies showed that *Shigella* can successfully invade epithelial cells derived from both small and large intestines (11). Based on the discrepancies between *in vivo* and *in vitro* observations, we hypothesize that differences between the luminal environments of small intestine and colon account for the preferential tropism of *Shigella* for the colon.

Bile is one important luminal factor that enteric pathogens must navigate in order to colonize and invade into the intestinal wall. It is a heterogeneous mixture of primary and secondary bile salts, cholesterol, phospholipids, and bilirubin that facilitates the digestion of fat in the small intestine (12). Pathogenic bacteria, including *Escherichia coli, Vibrio cholerae, Campylobacter jejuni* and *Shigella*, have evolved mechanisms to resist the antimicrobial properties of bile or even utilize bile as a signal to regulate virulence (13). Relevant here, upon exposure to bile salts, *Shigella* increases secretion of virulence factors, thus increasing adhesion to and invasion of host cells (14, 15). Additionally, long-term bile salt exposure *in vitro* induces *Shigella* biofilm formation, a phenomenon also observed when *V. cholerae* encounters bile salts (15, 16). A large percentage of bile salts are absorbed in the ileum, and approximately 400-800 mg of bile salts enter the colon daily (17). Thus, despite its importance in increasing pathogenic potency, it is unlikely that bile salts are the sole factor explaining the preferential colonic invasions by *Shigella*.

Concurrent with the inflow of bile into the small intestine, however, is the influx of bicarbonate (HCO_3_^-^), the physiological base used to buffer acidic contents from the stomach which creates a basic luminal environment in the distal small intestine (mean of 7.7, range for most subjects is pH 7.4-8) (18, 19). In contrast, the colonic lumen supports a more acidic environment (average pH is 6.4 though there is high variation, range of pH 5-8) (18, 19). Interestingly, bicarbonate modulates virulence factors, toxin production, and biofilm formation in *V. cholera* (16, 20, 21). However, there is a paucity of data regarding interactions between bicarbonate and *Shigella.* Given the drastic pH alteration from ileum to colon and the role of pH in regulating pathogenic activity of *Vibrio Cholerae*, we hypothesized that pH plays a vital role in regulating bile salts-dependent virulence of *Shigella* and its preferential pathogenesis in the colon.

In this study, we utilized previously established biofilm formation methods to test whether deoxycholate-induced biofilms of *Shigella flexneri* could form under various pH conditions. We demonstrated that more basic pH levels, as observed in the small intestine, attenuated biofilm formation without causing bacterial cell death. Additionally, we showed that basic pH conditions dispersed mature biofilms. RNA sequencing of *S. flexneri* under various pH conditions showed differences in transcriptional profiles of bacteria grown in deoxycholate with or without NaOH. Using a type III secretion system reporter strain of *Shigella*, we demonstrated that increasing pH could downregulate virulence as well. Collectively, these studies demonstrate that basic conditions, as found in the lumen of the small intestine, are not favorable for *S. flexneri* pathogenesis and provide new mechanistic insight into shigellosis pathogenesis.

## Results

### Basic Conditions Attenuate Deoxycholate-induced Biofilm Formation

Deoxycholate is a secondary bile salt that is present throughout both small and large intestines and is known to enhance secretion of *S. flexneri* virulence factors in planktonic culture (14). We initially tested if this bile acid affected the ability of *S. flexneri* to form biofilms in vitro (**Figure 1A**). To optimize conditions to produce *S. flexneri* biofilms, we grew *S. flexneri* in tryptic soy broth (TSB) supplemented with varying concentrations of sodium deoxycholate (NaDCA) and then measured the opacity of biofilms by optical density at 600 nm (OD_600_). Incubation of *S. flexneri* with 0.05% NaDCA stimulated formation of biofilms that approached 50% of the maximal opacity, and was thus suitable for testing the effects of external factors on biofilm formation (**Figure 1B**).

**Figure 1:**
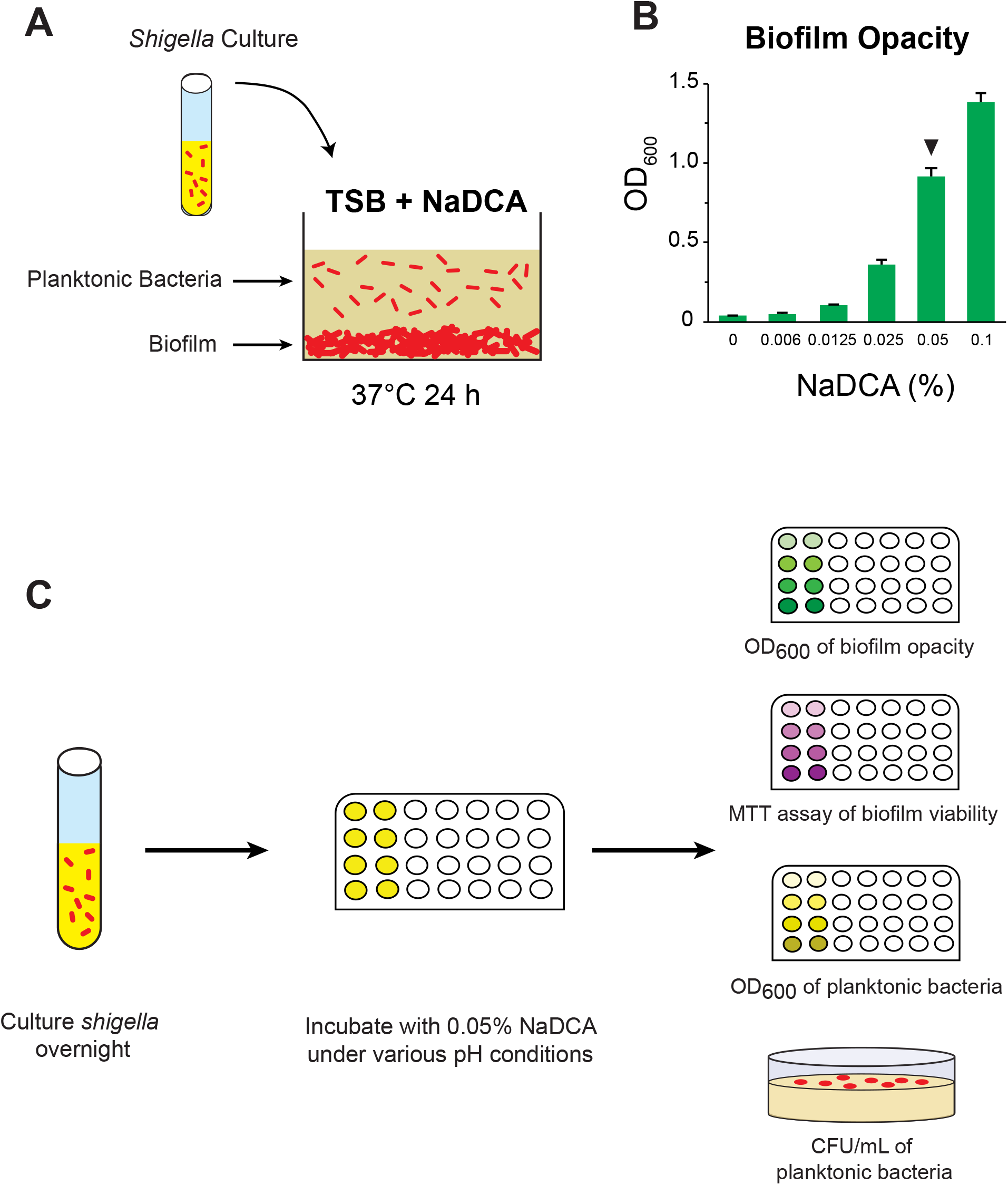
Experimental procedure utilized in this study. (A) Diagram showing the two resultant phases of bacterial growth after incubation in TSB with NaDCA overnight at 37°C. (B) The opacity of *Shigella* biofilms induced by increased percentages of sodium deoxycholate (NaDCA) for 24 h at 37°C. Note that 0.05% NaDCA induced biofilms that were approximately 50% of the maximal opacity and were selected for subsequent experiments (arrowhead). (C) Schematic of *Shigella* biofilm assays grown in 0.05% NaDCA and exposed to varying pH conditions. Biofilms were assayed for biofilm density (OD_600_ biofilm), viability (MTT), planktonic density (OD_600_ planktonic), and bacteria count (CFU/mL). NaDCA; sodium deoxycholate. TSB; tryptic soy broth.

We hypothesized that alterations of extracellular pH would impact biofilm formation, as the pH of the intestinal lumen is known to vary along its length (18). To simulate the basic conditions found in the small intestine, we adjusted the pH of the TSB media containing 0.05% NaDCA with increased concentrations of NaOH. We characterized the resulting biofilms by OD_600_ to measure opacity, MTT assay to assess microbial viability, and colony forming units to quantify the number of planktonic bacteria in the media that were not incorporated into the biofilm (**Figure 1C, 2A and 2B**). Increasing basic concentration in the TSB media decreased the opacity of the biofilms (**Figure 2C**) in a dose dependent manner with an almost complete loss of biofilm at 20mM NaOH (**Figure 2E**). Importantly, the pH that corresponds to 5 and 10mM is in the range of most human subjects in the distal small intestine (19). MTT assays, which have been utilized by many other biofilm studies, corroborated the reduction in biofilm formation with NaOH treatment (**Figure 2D**). Importantly, the attenuation of biofilm formation by increased pH did not appear to be secondary to enhanced bacterial killing by NaOH. In fact, the planktonic phase of *S. flexneri* that was maintained apically of the biofilm was enhanced by increased pH as measured by OD_600_ and CFU/mL (**Figure 2F and S1A**). Our interpretation is that there was an increased influx of bacteria into the planktonic phase as the biofilm was disassembled by basic conditions.

**Figure 2:**
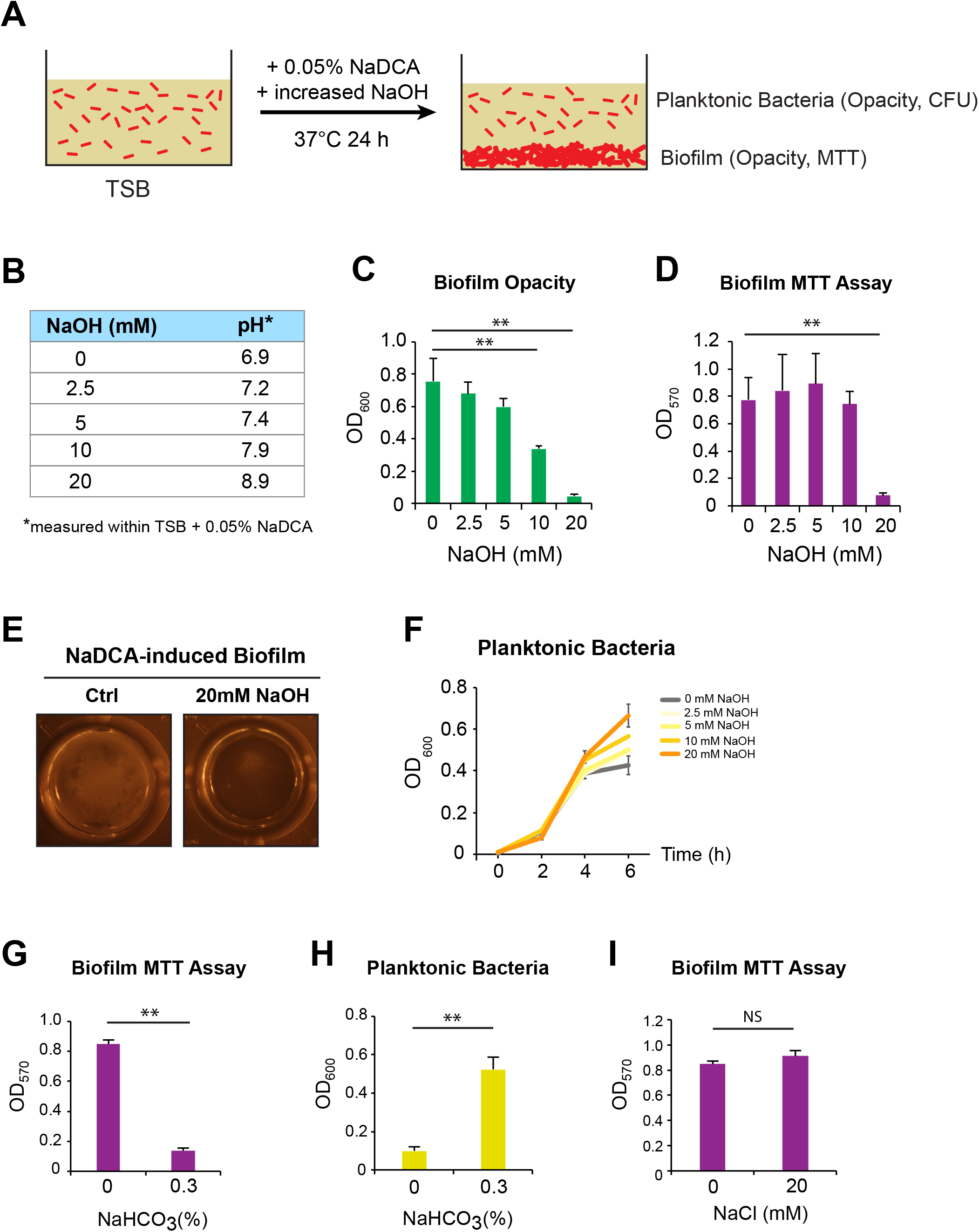
The effects of increased pH on formation of deoxycholate-induced biofilms. (A) Diagram describing the experimental steps for testing the effects of NaOH on biofilm formation induced by NaDCA. (B) A table showing the corresponding pH values when increased concentrations of NaOH were added to the TSB media with 0.05% NaDCA. (C-D) The opacity (as measured by OD_600_, C) and viability (as measured by MTT assay, D) of biofilms induced by 0.05% NaDCA with increased concentrations of NaOH after 24 h at 37°C. (E) Representative images of biofilms induced by NaDCA with or without 20mM NaOH. (F) Growth curve of the planktonic *Shigella* (measured by OD_600_) growing on top of the NaDCA-induced biofilms with increased concentrations of NaOH. (G-H) Biofilm viability (G) and planktonic growth curve (H) with or without 0.3% NaHCO_3_. (I) Biofilm viability with or without 20mM NaCl (as a control for ionic strength). n = 3 independent experiments. All bar graphs were plotted as Mean ± SD. Statistical significance was determined by Student’s t-tests with p <0.05. **=significant with two-tailed Student’s t-test.

Taken together, these data support the hypothesis that a basic environmental pH attenuates NaDCA-induced biofilm formation in *S. flexneri*, and this effect occurs at a range of pHs that is similar to the small intestinal lumen (**Figure 2B**). To further test that basic environments relevant to the small intestine can play a role in decreasing deoxycholate-induced biofilm formation, we repeated our assay with NaHCO3, the physiological agent responsible for modulating pH in the small intestine, and confirmed that a physiologically relevant bicarbonate concentration (0.3%) (22) could prevent biofilm formation as well (**Figure 2G and 2H**). To control for the possible effects induced by changes in ionic strength, we tested NaCl of the same concentrations and confirmed that NaCl did not affect biofilm formation (**Figure 2I**).

### Acidic Conditions Do Not Affect Biofilm Formation, but Decreases Viability of *S. flexneri*

To test if the acidic pH that *S. flexneri* encounters in the large intestine can influence biofilm formation, we supplemented TSB media with increasing concentrations of HCl (**Figure S1B**). In contrast to the increased biofilm density observed in basic conditions, we found that acidic pH did not significantly change the opacity of the biofilms (**Figure S1C**). Despite the presence of biofilms in all acidic conditions, the MTT assay showed diminished activity indicating that either the metabolism or number of bacteria surviving in the biofilms was decreased as the concentration of HCl was increased (**Figure S1D**). The number of planktonic bacteria growing in the media above the biofilm was also decreased with HCl addition as measured by CFU/mL (**Figure S1E and S1F**). Taken together, we showed that there was no reduction in biofilm assembly and formation in acidic conditions, despite decreased viability and/or metabolism for *S. flexneri*. These results support a model that *S. flexneri* biofilms are better supported in the colonic environment where there is an acidic milieu that does not prevent bile salt-induced biofilm formation.

### Environmental pH Modulates Dispersal of Mature *S. flexneri* Biofilms

The preceding experiments established that basic conditions prevent the assembly of biofilms from planktonic cultures. We next tested whether basic conditions affect biofilms that are already assembled and matured. First, we induced biofilm formation with NaDCA and allowed the biofilm to mature for 24 h, and then exchanged the growth media to TSB without NaDCA at a range of NaOH concentrations (**Figure 3A**). We found that the addition of 2.5, 5, or 10 mM NaOH did not significantly affect the biofilm opacity, though there was a trend towards reduction. A concentration of 20 mM NaOH treatment fully dispersed the mature biofilm as indicated by decreased opacity (**Figure 3B**). The effect of basic pHs on biofilm dispersal was not a potent as its effect on formation. These results were further supported by MTT assay (**Figure 3C**). These data suggest that the disruptive effects of NaOH on *S. flexneri* biofilms was not caused by its deprotonation effect on deoxycholate, but rather by its direct influences on the bacteria. We found that acidic environments did not significantly disperse mature biofilms or reduce bacterial viability (**Figure S2A and S2B**).

**Figure 3:**
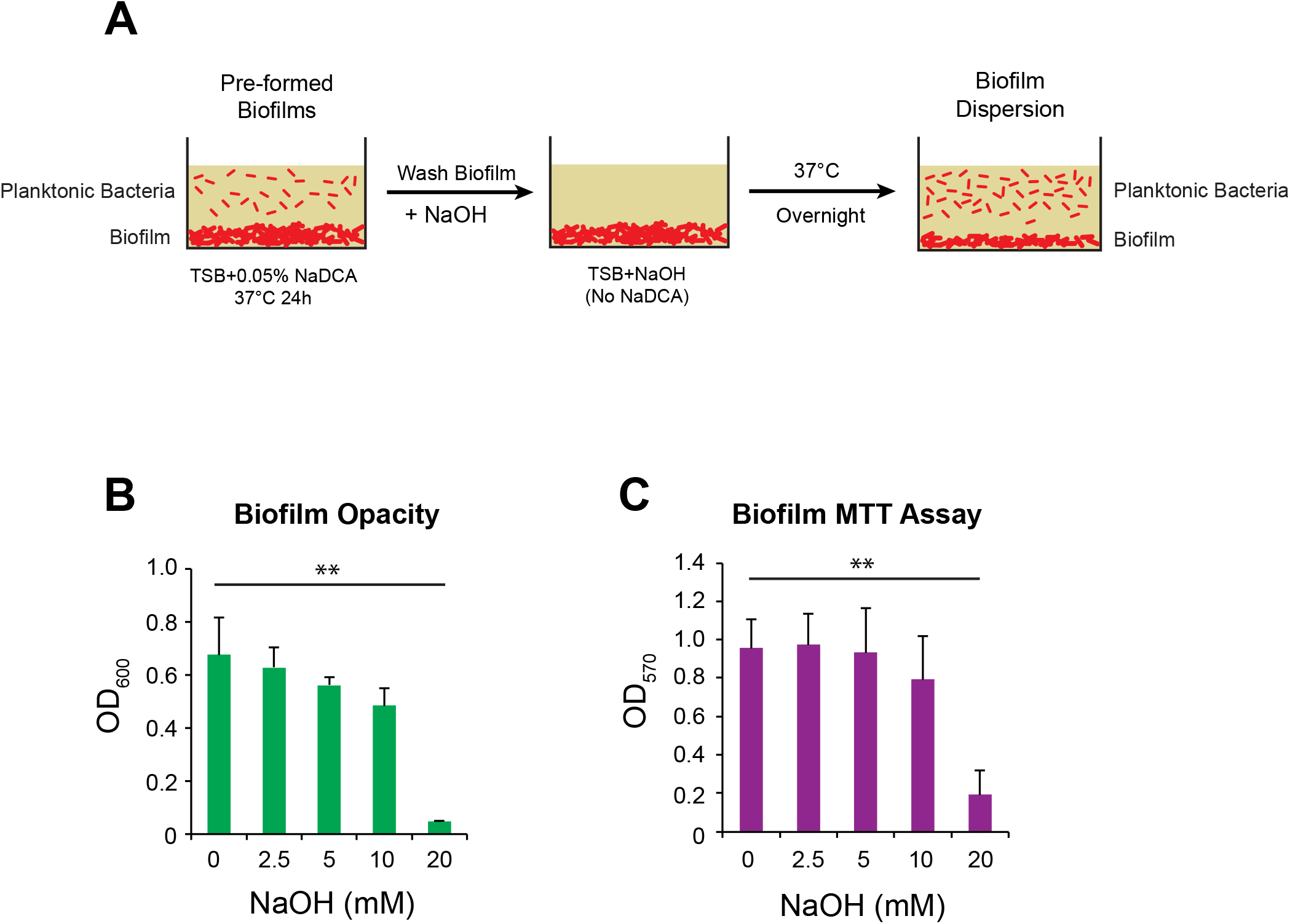
Dispersal of biofilms modulated by increased pH. (A) Illustration on the experimental procedures for testing the effects of NaOH on biofilm dispersion and disassembly. (B-C) The opacity (B) and viability (C) of pre-formed biofilms incubated with increased concentrations of NaOH for 24 h. n = 3 independent experiments. All bar graphs were plotted as Mean ± SD. Statistical significance was determined by Student’s t-tests with p <0.05. **=significant with two-tailed Student’s t-test.

### RNA-Sequencing Reveals that Basic Conditions Attenuate a Biofilm-Associated Transcriptomic Program

To gain additional insight into how pH affects *S. flexneri* biofilm formation, we performed RNA-sequencing to determine the transcriptomic alterations in *S. flexneri* caused by pH modulation. We performed this analysis using three experimental conditions that included (a) planktonic cultures in TSB only, (b) biofilms in TSB + 0.05% NaDCA, and (c) planktonic cultures in TSB + 0.05% NaDCA + 20 mM NaOH. A global view of the transcriptional profiles from these experimental groups, as shown by PCA plot, indicated that there were indeed transcriptional differences among each experimental group (**Figure S3A**).

To identify gene signatures that were associated with biofilm formation, we first compared the mRNA transcriptomic profiles between the deoxycholate-induced biofilm and planktonic *S. flexneri* grown in TSB only. Pathway analysis (DAVID 6.7) of genes upregulated in the biofilm showed an enrichment of ABC transporters, arginine metabolism, and sulfur metabolism (**Figure 4A**). Interestingly, arginine and cysteine metabolic pathways have been reported to be important in biofilm formation in other bacterial species (23, 24). We next compared the biofilm cultures grown with and without 20 mM NaOH. Pathway analysis revealed that the deoxycholate-induced biofilm gene enrichment of ABC transporters, arginine metabolism, and sulfur metabolism was repressed by the addition of NaOH (**Figure 4B).** Heatmap analysis of representative genes from these pathways further supported the specific increase in these genes only in the NaDCA conditions (**Figure 4C**). qPCR analysis on candidate genes that are involved in arginine and cysteine metabolism confirmed the RNA-seq results (**Figure S3B**). Taken together, these data demonstrated that basic conditions suppress a biofilm-associated gene program induced by deoxycholate in *S. flexneri*.

**Figure 4:**
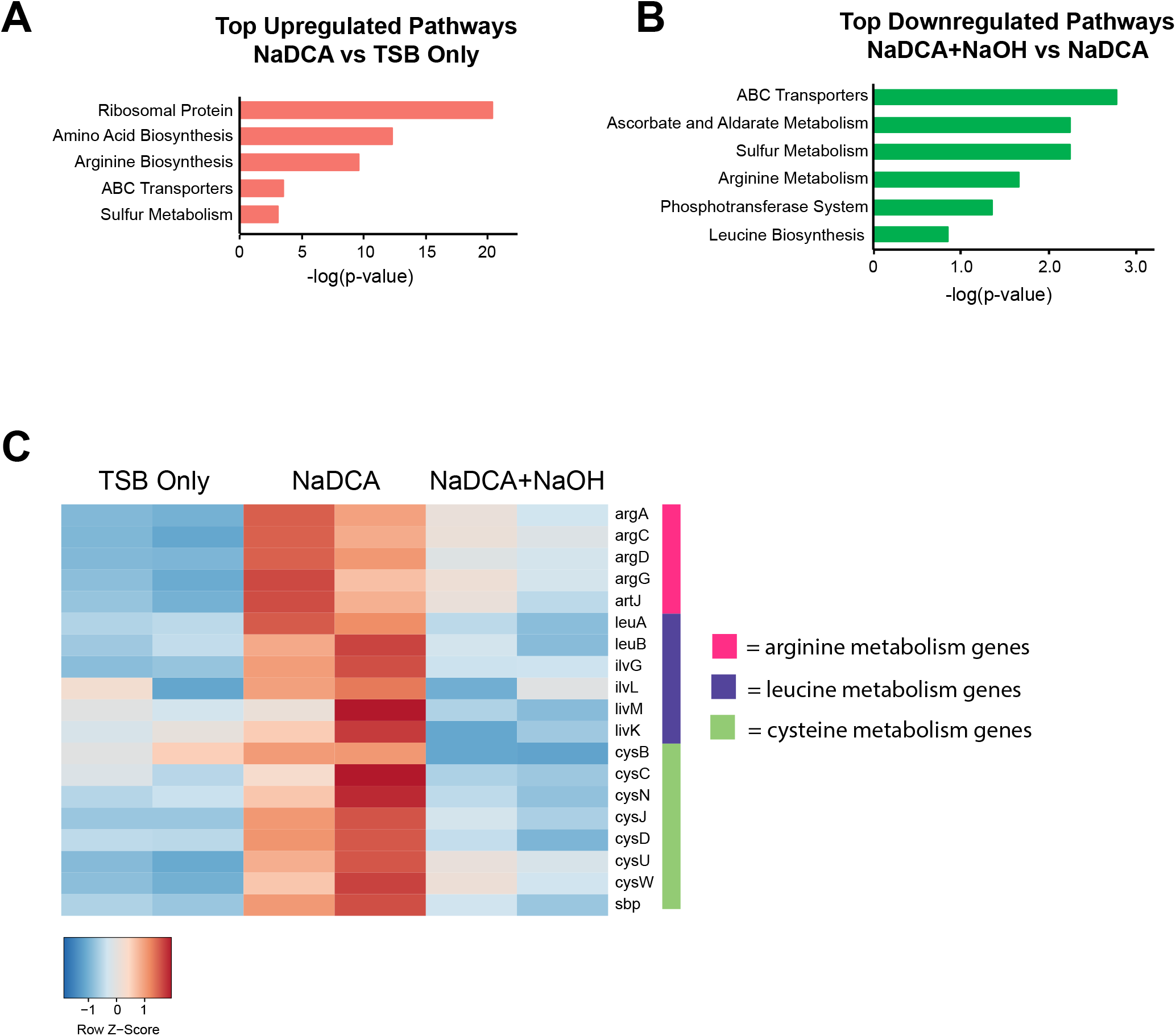
Transcriptomic analysis of the effects of increased pH on deoxycholate-induced biofilms. (A) Top upregulated pathways from *Shigella* cultured in TSB with 0.05% NaDCA compared to TSB only. (B) Top downregulated pathways from *Shigella* cultured in TSB with 0.05% NaDCA + NaOH compared to TSB with 0.05% NaDCA. (C) Heatmap showing the relative levels of genes involved in the arginine, leucine and cysteine metabolism pathways. Plots represented 2 independent experiments.

### Basic Conditions Prevent Induction of T3SS by Deoxycholate

In addition to stimulating biofilm formation, deoxycholate increases the expression of virulence factors in *S. flexneri*, especially those related to the type III secretion system (T3SS), the molecular machinery *S. flexneri* utilizes to invade into host cells (1). Given the attenuation of deoxycholate-induced biofilm formation in basic conditions, we hypothesized that addition of NaOH could also reduce virulence of *S. flexneri.* To test this, we utilized a previously constructed *Shigella* strain that contains a fluorescent reporter for the T3SS *(Shigella* M90T Sm pTSAR 2.4) (25). In this system, mCherry is constitutively expressed in all bacteria but GFP is expressed only when the T3SS apparatus is induced. We first demonstrated that addition of 0.2% NaDCA in TSB induced GFP production in this *Shigella* reporter strain, whereas bacteria growing in TSB alone did not fluoresce green. When the pH of the TSB was adjusted with increasing concentrations of NaOH, the T3SS activation by deoxycholate was reduced in a dosedependent manner with 5 and 10 mM concentrations of NaOH showing significant effects (**Figure 5A and 5B**). This experiment further supports the model that basic conditions deter *Shigella* from infection in the small intestine.

**Figure 5:**
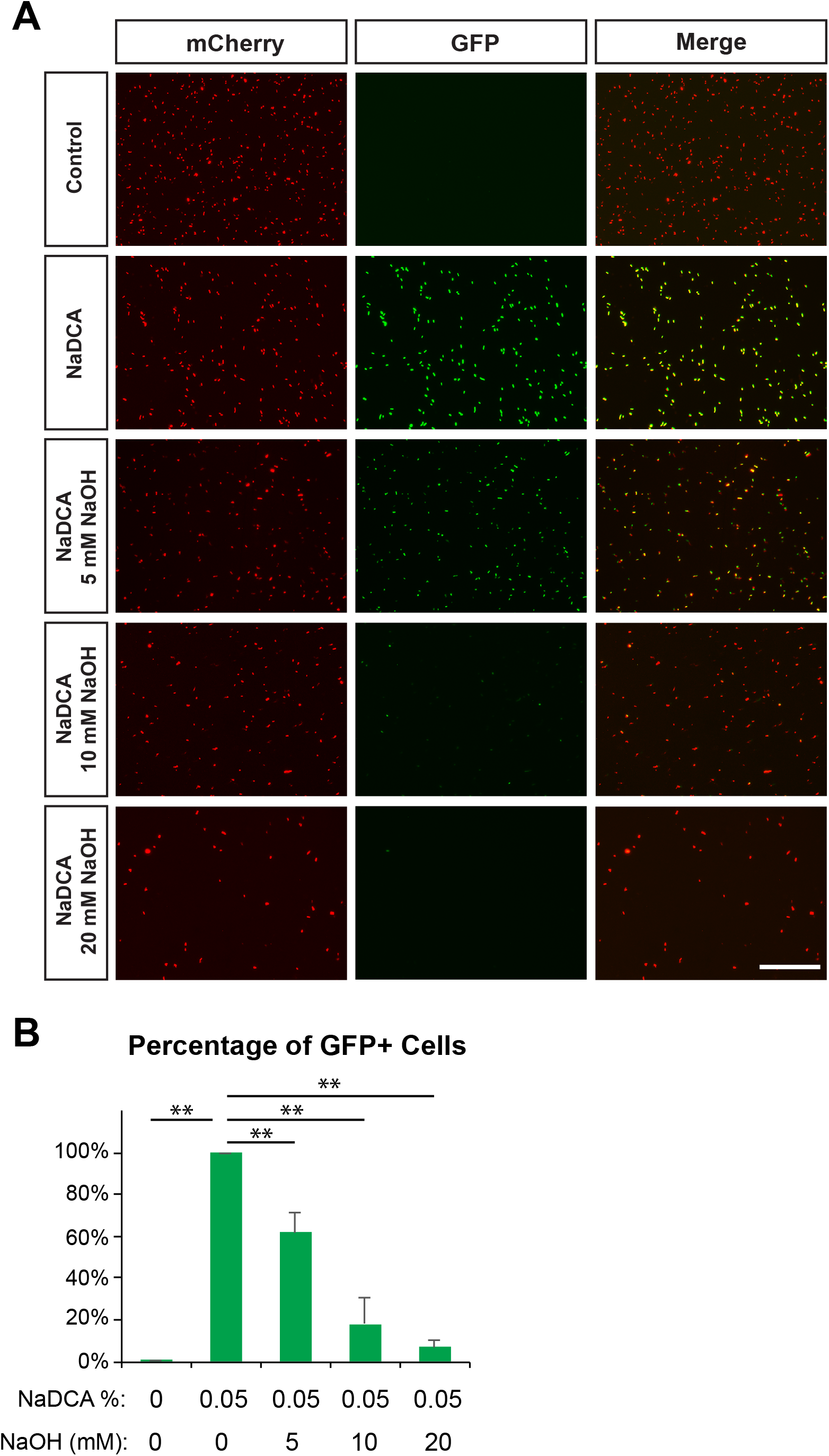
Increased pH inhibits deoxycholate-induced virulence of *Shigella*. (A) Images showing the effect of sodium deoxycholate and NaOH on a fluorescent reporter of *Shigella flexneri* which expresses GFP upon activation of the T3SS (with constitutive expression of mCherry as an internal control). Scale bar = 100μm. (B) Percentage of GFP+ cells out of mCherry+ cells under increased NaOH concentrations was plotted as Mean ± SD. n=2 independent experiments. Statistical significance was determined by Student’s t-tests. **p<0.01 with two-tailed Student’s t-test.

## Discussion

In this study, we utilized a series of *in vitro* bacterial assays, RNA-sequencing, and fluorescent microscopy to demonstrate that a decrease in deoxycholate-regulated biofilm formation and virulence induction of *S. flexneri* occurs under basic pH conditions. The results of these experiments support a model that *S. flexneri* would preferentially infect the colon over the distal small intestine. We propose that as *S. flexneri* transits through the distal small intestine, the basic pH of the luminal environment would act to reduce biofilm formation, virulence, and epithelial invasion. As it reaches the more acidic colonic lumen, the bacteria could increase biofilm formation and virulence-related gene expression which would facilitate epithelial invasion at this site.

*V. cholera*, a pathogen that preferentially colonizes the small intestine, serves as an important comparator to *Shigella*. In both bacteria, bile salts can induce biofilm formation (16). However, bicarbonate upregulates virulence factors and toxin production in *V. cholera*, whereas the same condition reduces virulence in *Shigella* (13, 15). Thus, our study of *Shigella* and previous studies of *V. cholera* highlight the importance of considering environmental factors in the lumen when studying pathogenesis of specific enteric pathogens that infect different regions of the gut.

Recent advances in the detection of intestinal pathogens using qPCR revealed that *Shigella* may be twice as prevalent as previously estimated (26). Due to the lack of available vaccines for shigellosis, antibiotics are currently the first line of treatment for this diarrheal disorder (27). However, antibiotic resistance has become an increasingly severe problem in part exacerbated by the excessive antibiotic use. Thus, we urgently need to find alternative approaches to treat bacterial diseases like shigellosis (28). In this study, we elucidate a pH-dependent mechanism by which the small intestine evades *Shigella* invasion. With further investigations on specific pathways that mediate the reduction of virulence and biofilm formation under basic pH conditions, we can potentially find targets to therapeutically treat shigellosis.

Bacterial biofilms play a clear role in physiology and pathophysiology in many different biologic systems. For example, *Vibrio fischeri* produces biofilms in the light organ of the Hawaiian squid, *Pseudomonas aeuginosa* produces biofilms in the lung airways that that can lead to diminished respiratory function and polymicrobial biofilms on the enamel surface of teeth. (23, 24, 29). Despite the extensive number of studies on bacterial biofilms, there is been minimal characterization of deoxycholate-induced *Shigella* biofilms. Through RNA-sequencing of *Shigella* biofilms, we showed that the arginine and cysteine metabolism pathways were upregulated and enriched. The arginine and cysteine metabolism pathways are also associated with *P. aeruginosa* biofilms and *V. fischeri* biofilms, respectively (30, 31). Because biofilms in the gut have been potentially associated with colorectal cancer, inflammatory bowel disease, and various enteric infections, our studies may help further the investigation of biofilms related to disorders of the gastrointestinal tract (32, 33).

Although more work is needed to further elucidate the pathways and host factors that control *Shigella* virulence and biofilm formation, our work expands on our knowledge of *Shigella* pathogenesis and provides a unique perspective into studying mechanisms of infection. This will prove to be imperative as we are slowly attempting to uncover more and more interactions between the host and pathogen.

## Materials and Methods

### Biofilm Formation Assays

*Shigella flexneri* strain 2a 2457T was purchased from ATCC (#700930) and used within 6 months of receipt. *S. flexneri* was inoculated into LB media from the original commercial stock and grown 20h at 37°C with shaking at 250rpm. After this incubation, the bacteria were pelleted by centrifugation, resuspended in tryptic soy broth (TSB), and measured by OD_600_. The bacterial suspension was then diluted with TSB and adjusted to an OD_600_ of 0.5. From this stock, the bacteria were further diluted 1:800 in TSB that included 0.05% NaDCA. The bacterial suspension was then added to a 96-well plate (180 μL/well). For experiments 20 μL of H_2_O, diluted HCl, or diluted NaOH was added to each well to achieve desired final concentrations. The plate was incubated stationary at 37°C for 24h to allow for biofilm formation. The supernatant above the resultant biofilm was removed and set aside for OD_600_ or CFU/mL measurement; the biofilms were washed with 100 μL H_2_O twice and measured by OD_600_ before the MTT solution (100 μL; 0.5 mg/mL in TSB) was added to the biofilms. The plate was incubated at 37°C for 10 minutes before the MTT solution was removed and dissolved with 100 μL DMSO. The OD_570_ of the resulting solution was measured by Cytation 5. The planktonic cells in the supernatant were sequentially diluted in PBS and plated onto LB plates for CFU/mL quantification. A similar procedure was utilized for the testing of NaHCO_3_, NaCl, and the dose curve of NaDCA.

### *S. flexneri* Growth Curve

*S. flexneri* was grown overnight in LB media. The bacteria were collected with centrifugation, adjusted to OD_600_ of 0.5, and diluted 1:20 in TSB. The bacterial suspension was added to 96-well plates (180 μL/well) and then additional H_2_O, HCl, or NaOH solutions were added to achieve final volume of 200 μL and desired concentrations. The plate was incubated at 37°C with shaking, and the OD_600_ of the plate was measured at 2, 4 and 6 h.

### Biofilm Dispersion Assays

An overnight culture of *S. flexneri* in LB was adjusted to OD_600_ of 0.5 and further diluted 1:800 in TSB with 0.05% NaDCA. The diluted bacterial suspension was added to 96-well plates (200 μL/well) and incubated without shaking at 37°C for 24 h. After 24 h, the supernatant above the biofilm was removed, the biofilms were washed once with H_2_0 (100 μL) and a new solution of TSB with various concentrations of NaOH or HCl (200 μL/well) was added on top of the biofilms. After incubation at 37°C with no shaking for 24 h, the resulting biofilms were measured by OD_600_ and the MTT assay as described above.

### RNA Sequencing of *S. flexneri*

The diluted solution of bacteria from an overnight culture (first adjusted to OD_600_ of 0.5, then 1:20 dilution in TSB) was further diluted to 1:800 in TSB +/− 0.05% NaDCA. For TSB only control samples, an aliquot of 200uL/well of the above dilutions (without NaDCA) was added to into 96 well plates. To assess the effects of NaDCA and/or NaOH, an aliquot of 180 μL/well of the above dilutions (with NaDCA) was added with 20 μL of either H_2_O or 200mM NaOH. The plates were incubated stationary at 37°C for 24h. The planktonic bacteria in the wells containing TSB only (no NaDCA) or TSB with 0.05% NaDCA and 20mM NaOH were collected and pelleted with centrifugation (15 min, 4C) and lysed according to instructions of RiboPure RNA Purification kit for bacteria (Invitrogen). Biofilms from wells containing TSB and 0.05% NaDCA were washed and directly lysed with the same kit. RNA was then purified using the RNeasy Mini Kit (Qiagen). Genomic DNA removal was performed according to instructions of TURBO DNA-free Kit (Invitrogen).

Libraries were prepared according the manufacture protocol and sequenced with Illumnia HiSeq3000. The Ensembl release 76 top-level assembly STAR version 2.0.4b was used to align RNA-seq reads and the Subread:feature Count version 1.4.5 was used to derive gene counts. Gene counts were then imported into and analyzed with R/Bioconductor package EdgeR^5^ for normalization and then imported into R/Bioconductor package Limma. Differential expression was determined with Benjamini-Hochberg false-discovery rate adjusted p-values less than or equal to 0.05. PCA plot was made with DDSeq2 package based on all gene counts. Pathway analysis was performed by utilizing the Database for Annotation, Visualization and Integrated Discovery (DAVID) v6.7 (https://david-d.ncifcrf.gov).

For qPCR verification, 500ng of RNA was converted to cDNA using iScript Reverse Transcription Supermix Kit (Bio-Rad) and qPCR was performed with TB Green Advantage qPCR Premix. RecA was used as a housekeeping gene. Primers utilized are shown in **Table S1**.

### Fluorescence Microscopy to Probe Activation of T3SS

A Congo red positive colony of *Shigella* M90T Sm transformed with the pTSAR 2.4 plasmid (a kind gift from Dr. François-Xavier Campbell-Valois) was grown in TSB overnight. The strain was transformed following protocol as previously reported (25). The overnight culture was diluted 1:100 into TSB solutions with or without 0.2% NaDCA under various pH conditions (adjusted with H_2_O, HCl, or NaOH), and incubated for 3h without shaking at 37°C. The activation of type III secretion system was assessed with the GFP channel by fluorescent microscopy (25).

## Achknowledgements

We thank Dr. François-Xavier Campbell-Valois for providing *Shigella* M90T Sm and pTSAR 2.4 plasmid. We also thank Shanshan Xiong for her help in this project. I.L.C is supported by a fellowship for MA/MD program from Washington University School of Medicine. Crohn’s & Colitis Foundation provided support.

## Supplemental Figure Legends

**Figure S1:**
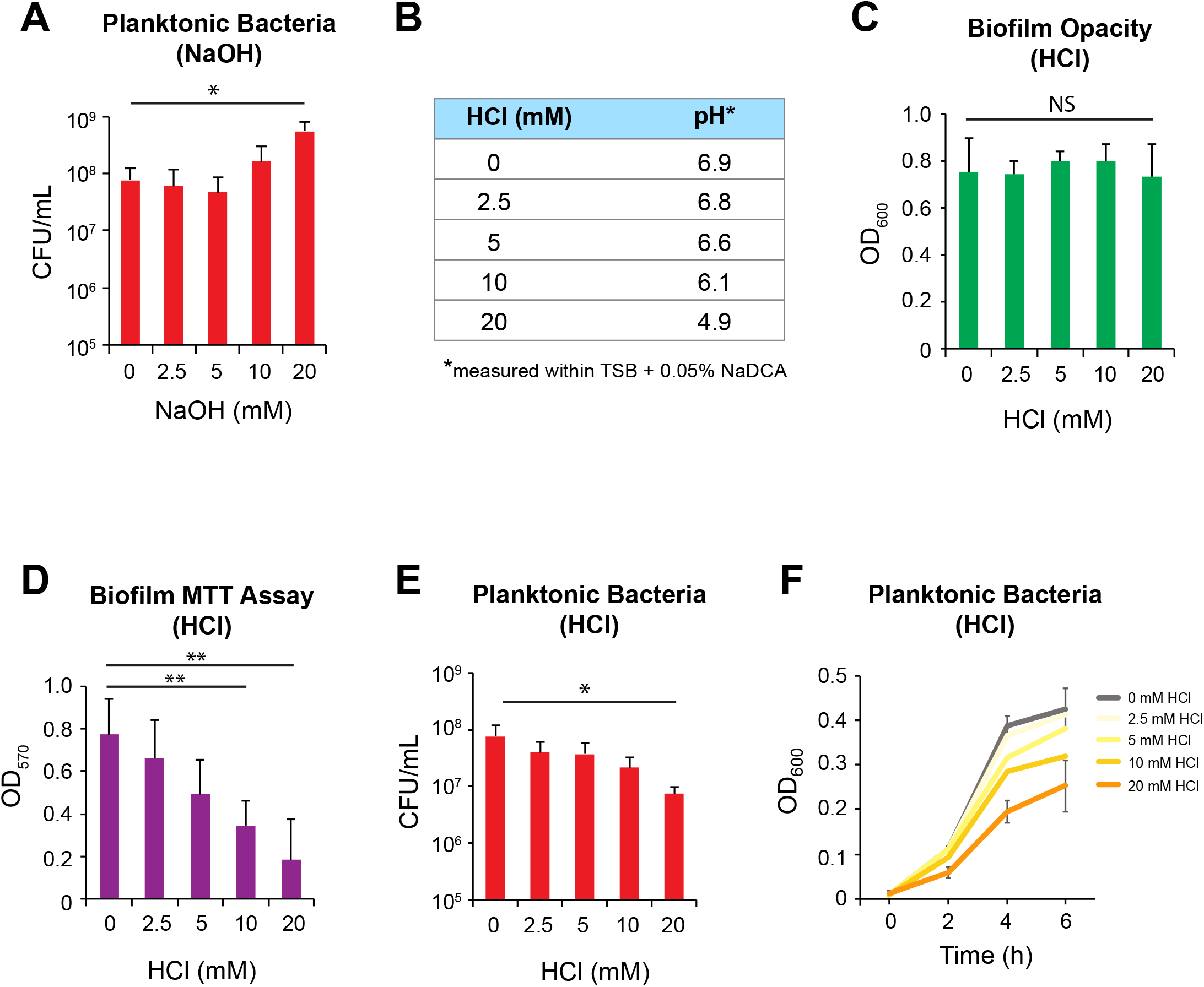
The effect of decreased pH on *Shigella* biofilm formation. (A) CFU/mL measurement of the planktonic *Shigella* growing on top of NaDCA-induced biofilms with increased concentrations of NaOH after 24 h at 37°C. (B) A table showing the corresponding pH values when increased concentrations of HCl were added to the TSB media with 0.05% NaDCA. (C-D) The opacity (C) and viability (D) of biofilms induced by 0.05% NaDCA with increased concentrations of HCl. (E) CFU/mL measurement of the planktonic *Shigella* growing on top of NaDCA-induced biofilms with increased concentrations of HCl after 24 h at 37°C. (F) Growth curve of the planktonic *Shigella* (measured by OD_600_) growing on top of the NaDCA-induced biofilms with increased concentrations of HCl at indicated time points. n = 3 independent experiments. All bar graphs were plotted as Mean ± SD. Statistical significance was determined by Student’s t-tests with p <0.05. **=significant with two-tailed Student’s t-test. *=significant with one-tailed Student’s t-test. NS=not significant.

**Figure S2:**
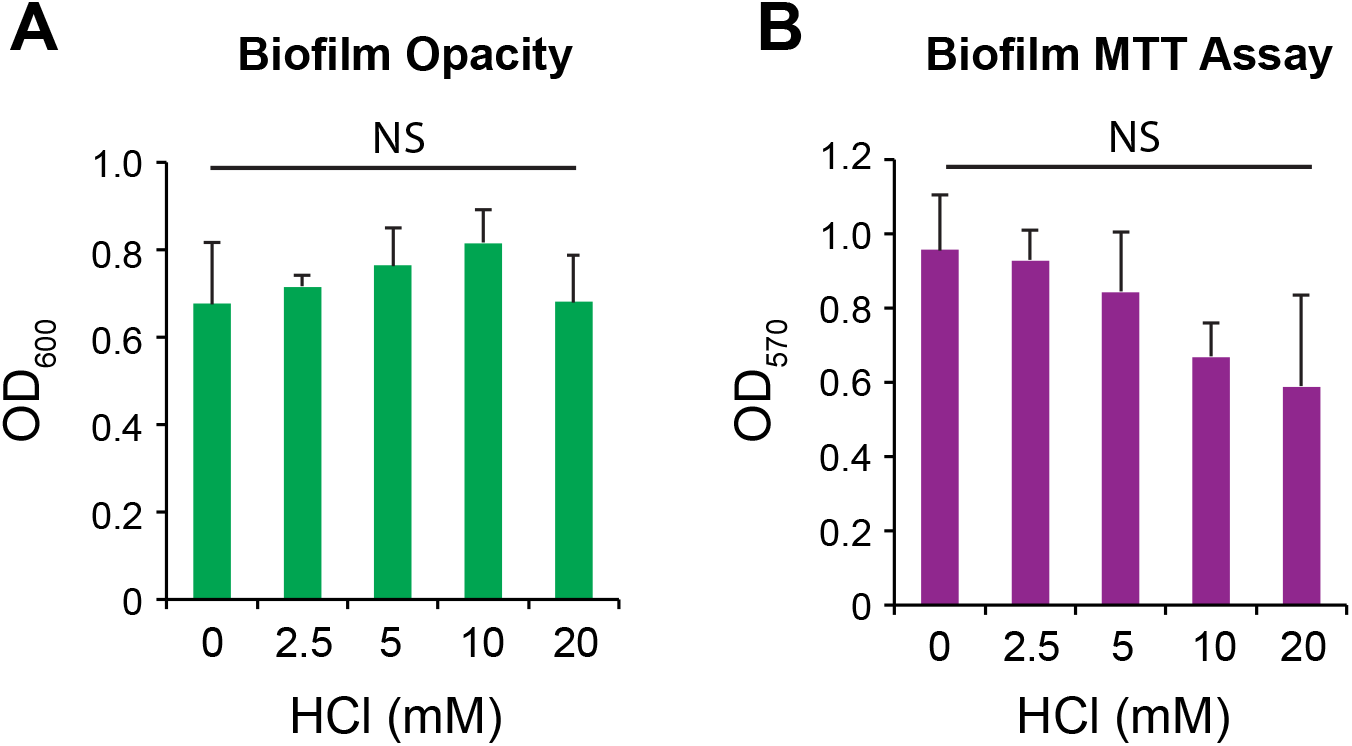
Dispersal of biofilms modulated by decreased pH. (A-B) The opacity (A) and viability (B) of pre-formed biofilms incubated with increased concentrations of HCl for 24 h. n = 3 independent experiments. All bar graphs were plotted as Mean ± SD. Statistical significance was determined by Student’s t-tests with p <0.05. NS=not significant.

**Figure S3:**
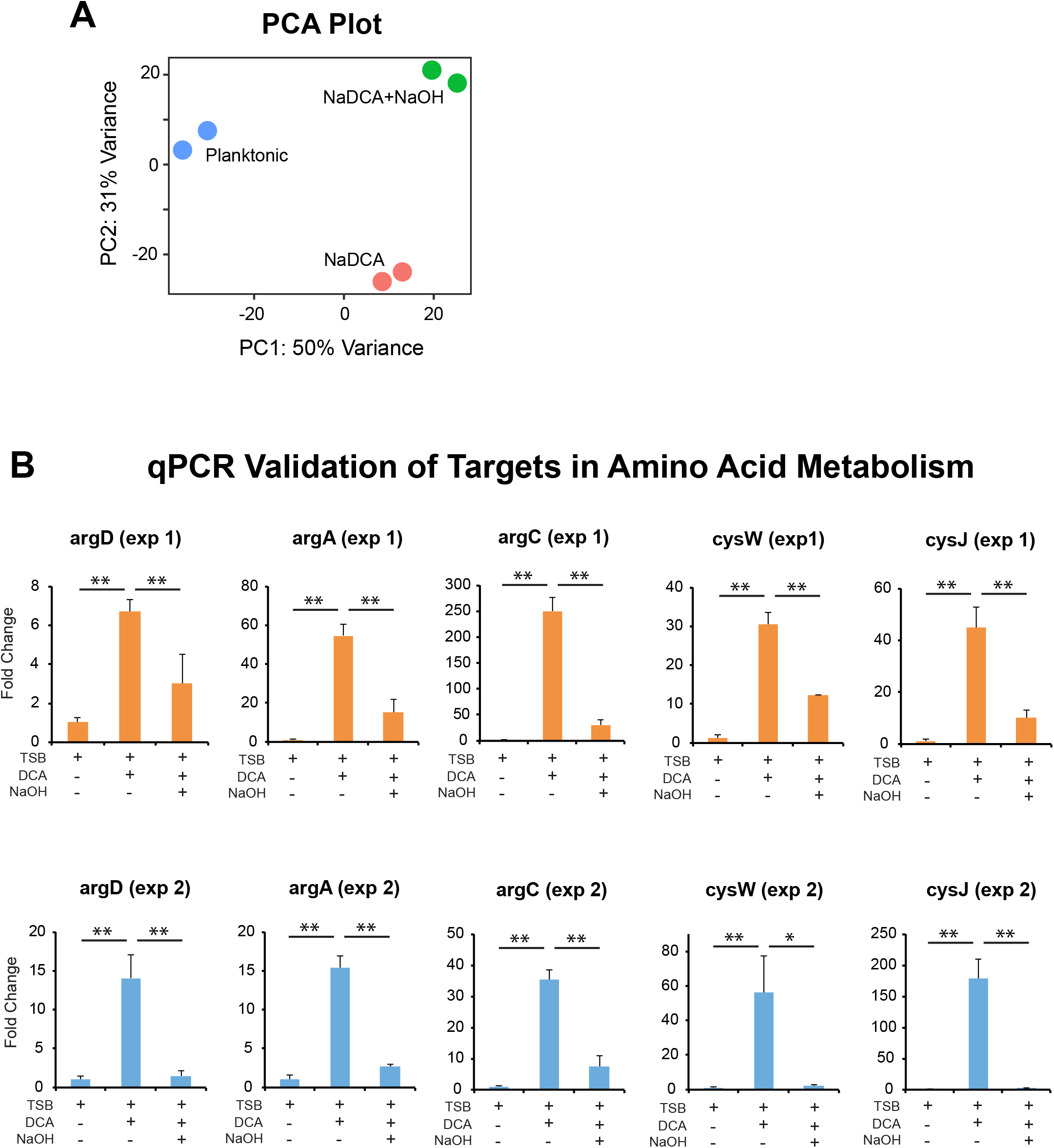
Verification of RNA-sequencing results with qPCR. (A) PCA plot of the transcriptomes from *Shigella* cultured in TSB only, TSB with 0.05% NaDCA and TSB with 0.05% NaDCA + 20mM NaOH. (B) qPCR results of various genes involved in arginine and cysteine metabolism were plotted as Mean ± SD for 2 independent experiments. n=3 technical replicates for each experiments. Statistical significance was determined by Student’s t-tests. **p<0.01 with two-tailed Student’s t-test. *p<0.05 with two-tailed Student’s t-test.

