## Supplemental Material for "Biofilm Formation and Virulence of *Shigella flexneri* is Modulated by pH of Gastrointestinal Tract"

| Gene name | | Primers 5'-> 3' |
|---|---|---|
| ansB | Forward Primer | TACCATTACCCACACCAGCG |
| | Reverse Primer | AGCGATACGCCATTCGATGT |
| asnA | Forward Primer | GCATATCGGCCAGGTTCAGT |
| | Reverse Primer | TTACAGCAGAGAAGGGACGC |
| napA | Forward Primer | CTTTCACTTTGTCGCCACGG |
| | Reverse Primer | AGTACGACCTGTGGCTCTCT |
| ccmF | Forward Primer | CCCATAGCGGATACACGGAC |
| | Reverse Primer | CTGTGCTTGGCGTTAGGGAT |
| ccmE | Forward Primer | GCTTTCTCAACTTCTGGCGG |
| | Reverse Primer | GCGAGCTGGAAAAAGGCAAT |
| cysW | Forward Primer | ACGGTGTTGTAGTCCTGCTC |
| | Reverse Primer | ACTTTCGCTGCCGTTACAGA |
| cysB | Forward Primer | GCGTATCGTTTTCACGGCAA |
| | Reverse Primer | CATGCTGGCAATGACCCCTA |
| cysJ | Forward Primer | ACTTCGCCCTCTTCTTCCAC |
| | Reverse Primer | ACCCCGCGTCTCTATTCCAT |
| argD | Forward Primer | CGGCATTCAGCACCATTACG |
| | Reverse Primer | TAATTGGCGCGGAGCTGAAA |
| argA | Forward Primer | AGTCTCAATAACACGCCGCT |
| | Reverse Primer | ATTGCACCGCTTTCCAGTTG |
| argC | Forward Primer | TGACTGTTTCAGCGCAAAGC |
| | Reverse Primer | GGCGAGAAACACTACGTCCA |
| recA | Forward Primer | CAGGCAGTTGCATTCGCTTT |
| | Reverse Primer | TCTACGGCGAACTGGTTGAC |
